# Pan-cancer analysis of the metabolic reaction network

**DOI:** 10.1101/050187

**Authors:** F. Gatto, J. Nielsen

## Abstract

Metabolic reprogramming is considered a hallmark of malignant transformation. However, it is not clear whether the network of metabolic reactions expressed by cancers of different origin differ from each other nor from normal human tissues. In this study, we reconstructed functional and connected genome-scale metabolic models for 917 primary tumors based on the probability of expression for 3,765 reference metabolic genes in the sample. This network-centric approach revealed that tumor metabolic networks are largely similar in terms of accounted reactions, despite diversity in the expression of the associated genes. On average, each network contained 4,721 reactions, of which 74% were core reactions (present in >95% of all models). Whilst 99.3% of the core reactions were classified as housekeeping also in normal tissues, we identified reactions catalyzed by *ARG2, RHAG, SLC6* and *SLC16* family gene members, and *PTGS1* and *PTGS2* as core exclusively in cancer. The remaining 26% of the reactions were contextual reactions. Their inclusion was dependent in one case (*GLS2*) on the absence of *TP53* mutations and in 94.6% of cases on differences in cancer types. This dependency largely resembled differences in expression patterns in the corresponding normal tissues, with some exceptions like the presence of the *NANP*-encoded reaction in tumors not from the female reproductive system or of the *SLC5A9-*encoded reaction in kidney-pancreatic-colorectal tumors. In conclusion, tumors expressed a metabolic network virtually overlapping the matched normal tissues, raising the possibility that metabolic reprogramming simply reflects cancer cell plasticity to adapt to varying conditions thanks to redundancy and complexity of the underlying metabolic networks. At the same time, the here uncovered exceptions represent a resource to identify selective liabilities of tumor metabolism.

## INTRODUCTION

Dysregulation of cellular metabolism has been implicated in the progression of several cancers as a consequence of oncogenic mutations (*1*, *2*). Despite the fact that the regulatory programs underlying the observed metabolic shifts should be tumor specific, it is also known that at the system level metabolic regulation is substantially similar between the tumor and its tissue of origin (*3*-*5*). Even when shown to be selectively essential to cancer cells, the diversity of metabolic phenotypes associated with cancer questions the extent to which these regulatory programs are context-dependent rather than tumor-specific (*6*, *7*). For example, glucose metabolism was shown to vary within tumor regions and between human tumors in lung cancer patients (*8*) or to depend strongly on the initiating oncogenic mutation and the tumor tissue of origin in genetically engineered mice (*9*). Human metabolism is a highly complex system, accounting for thousands of reactions and metabolites that interact with the environment to form a connected and functional metabolic network (*10*). It is plausible that metabolic shifts so far associated with different cancers are yet another expression of the plasticity of these cells to ever-changing conditions in their genome and their environment (*11*), with the advantage that in metabolism this adaptation can leverage on the high redundancy and complexity of the human metabolic network. The aim of this study was therefore to characterize the landscape of metabolic reactions expressed in different cancers, to define their occurrence depending on the cancer type or mutations in key cancer genes, and finally to identify any difference from metabolic reactions normally expressed in human tissues.

Networks represent the natural structure of biological systems, including metabolism (*12*). Network-dependent analyses not only enhance the interpretability of genome-scale data by providing a context, but also highlight knowledge gaps and experimental artifacts, e.g. networks can have been used to build a systematic gene ontology, where manual curation would have been biased towards well-studied cellular processes (*13*). In cancer metabolism, network-based approaches have unveiled non-trivial metabolic dependencies of the studied cancers (*14*-*19*). In light of this, we sought to reconstruct the metabolic networks of 1,082 primary tumor samples in order to obtain a network-dependent landscape of metabolic reactions occurring in different cancers. These were reconstructed in the form of genome-scale metabolic models (GEMs) (*20*-*22*) in order to derive viable metabolic networks. GEMs encode all metabolic reactions performed by the gene products occurring in a sample, while ensuring that included reactions can carry flux during the simulation of essential metabolic tasks, such as synthesis of biomass constituents. Once the GEMs were reconstructed, we analyzed the underlying metabolic networks to address the above-defined goals of the study.

## RESULTS

### Reconstruction of 917 cancer genome-scale metabolic models

In order to explore the landscape of metabolic reactions in cancer, we reconstructed the metabolic network in each primary tumor of a cohort consisting of 1,082 patients, spanning 13 cancer types. Clinical, genetic, and gene expression data were retrieved for each sample from The Cancer Genome Atlas (TCGA). Each metabolic network was reconstructed in the form of a genome-scale metabolic model (GEM) (*21*), here-on referred to as model. The reconstruction was performed by mapping RNA-seq gene expression data from each tumor sample in a reference generic GEM of the human cell, *HMR2* (*23*), using the tINIT algorithm (*24*) and by estimating the probability that a gene is truly expressed in that sample using an *ad hoc* Bayesian statistical framework. As per algorithm formulation (*25*), each reconstructed GEM results in a connected and functional metabolic network. A connected GEM can carry flux in each reaction under standard medium conditions, and this requirement ensures for example that expressed genes that become isolated due to the malignant transformation are eliminated from the network. A functional GEM, on the other hand, can simulate a flux through 56 fundamental metabolic tasks, including biomass growth. Noteworthy, samples used in this study represent a mixed population of cells, predominantly constituted by cancer cells (>80% of tumor nuclei in each sample according to TCGA quality criteria). Therefore, each GEM should be thought as the metabolic network expressed in the tumor, rather than solely representing the cancerous cells.

We enforced quality criteria to the reconstructed GEMs to ensure to work with metabolic networks that include and exclude genes primarily because not likely to be expressed in that sample. Out of the initial 1,082 samples, 917 reconstructed GEMs passed these quality criteria (Fig. S1). We observed that many discarded models were derived from pancreatic adenocarcinoma (PAAD) and clear cell renal cell carcinoma (KIRC) samples. Specifically, GEMs for 15 PAAD samples (52%) and 53 KIRC samples (51%) were discarded because they failed to converge to an optimal solution (Fig. S2). The lack of convergence in the case of KIRC is consistent with the notion of a highly compromised metabolic network that we uncovered previously for this cancer type (*5*, *26*). Finally, we inspected whether the genes included in a GEM have indeed a higher expression level in the corresponding sample compared with the genes excluded. This was the case both when looking at individual models (Fig. S3B) and all models simultaneously (Fig. S3A). Nevertheless, we noted that a fraction of lowly expressed genes was still included in most reconstructed models, likely to preserve connectivity and functionality of the metabolic networks. Taken together, we were able to reconstruct GEMs representative of the metabolic network expressed in 917 cancers.

### Classification of metabolic reactions in core or contextual

Compared to the reference GEM, *HMR2*, which contains 8,184 reactions associated with 3,765 genes, the reconstructed GEMs featured on average 4,721 reactions and 1,995 genes (Fig. S4). We observed small variations in the number of reactions, but substantial differences in the number of genes included across GEMs. We performed principal component analysis (PCA) on the gene inclusion matrix, that is the binary *g* × *m* matrix where 1 indicates inclusion of *HMR2* gene *g* in the *m*-th GEM, and 0 vice versa. PCA revealed genetic diversity across GEMs attributable to the different cancer type (Fig. 1A), even by taking into account that the reconstruction might be biased in the case of PAAD and KIRC types. Nevertheless, PCA on the reaction inclusion matrix virtually abolishes the above-seen diversity among models (Fig. 1B). This suggest a considerable similarity in metabolic functions in cancer, in spite of gene expression heterogeneity.

**Figure 1.**
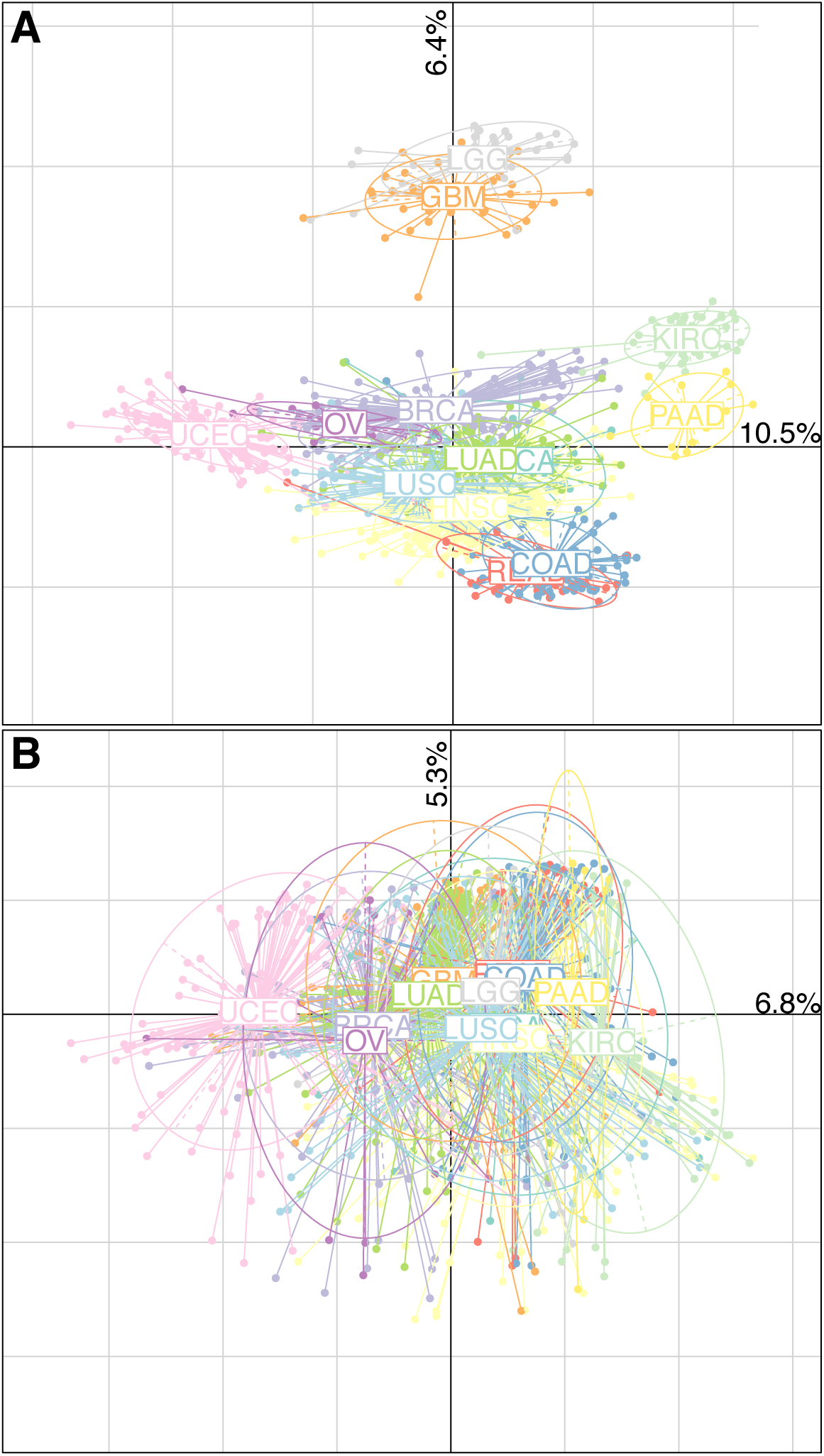
Principal component analysis of 917 cancer genome-scale metabolic models based on gene (A) and reaction (B) inclusion from the reference generic human model. Models are grouped by cancer type. Key: BLCA – Bladder adenocarcinoma, BRCA – Breast carcinoma, COAD – Colon adenocarcinoma, HNSC – Head and neck squamous cell carcinoma, GBM – Glioblastoma multiforme, KIRC – Clear cell renal cell carcinoma, LGG – Low grade glioma, LUAD – Lung adenocarcinoma, LUSC – Lung squamous cell carcinoma, OV – Ovarian carcinoma, PAAD – Pancreatic adenocarcinoma, READ - Rectum adenocarcinoma, UCEC – Uterine corpus endometrial carcinoma.

To shed light on the extent of the similarity in terms of reactions across cancers, we classified a reaction as “core” if included in >95% of all models, “absent” if excluded in >95% of all models, and “contextual” if otherwise (Fig. 2A-S5). At this threshold, 3,510 reactions are core (95% bootstrap confidence interval [CI], 3,421 to 3,599), 3,455 are contextual (95% CI, 3,367 to 3,539), and 1,219 are absent (95% CI, 1,157 to 1,284) (Fig.2B). Since reactions can be associated with more than one gene (i.e. isoenzymes), we further distinguished core reactions into “pan” if included in all models because of the same gene-reaction association or “iso” if otherwise (Fig. 2A). Finally, some reactions have no gene associations (e.g. spontaneous reactions), but they were classified as core because they connected other core reactions. We termed these “conn” reactions. Among core reactions, 2,850 are pan reactions (95% CI, 2,807 to 2,897), 69 are iso reactions (95% CI, 52 to 85), and 590 are conn reactions (95% CI, 546 to 632) (Fig. 2B). Considering an average of 4,721 reactions per GEM, this suggests that 74% of all metabolic reactions in the expressed network are found in any given cancer. Importantly, we calculated a significantly higher fraction of core reactions than expected by chance, using 1,000 sets of 917 randomly generated models, with equivalent gene inclusion diversity as in the reconstructed set, but no constrain on connectivity or functionality (Fig. 2B). A bird’s eye view on the generic KEGG metabolic map showed that the core reactions mostly cover primary metabolic pathways (e.g. energy, nucleotide and lipid metabolism), while contextual reactions appear more peripheral (e.g. glycan metabolism, Fig. 2C), which was confirmed when we grouped core vs. contextual reactions in *HMR2* metabolic subsystems (Fig. S6). Taken together, these results suggest that, despite heterogeneity in the expressed metabolic genes, distinct cancers express a strikingly similar metabolic network, which can support flux connectivity and functionality.

**Figure 2.**
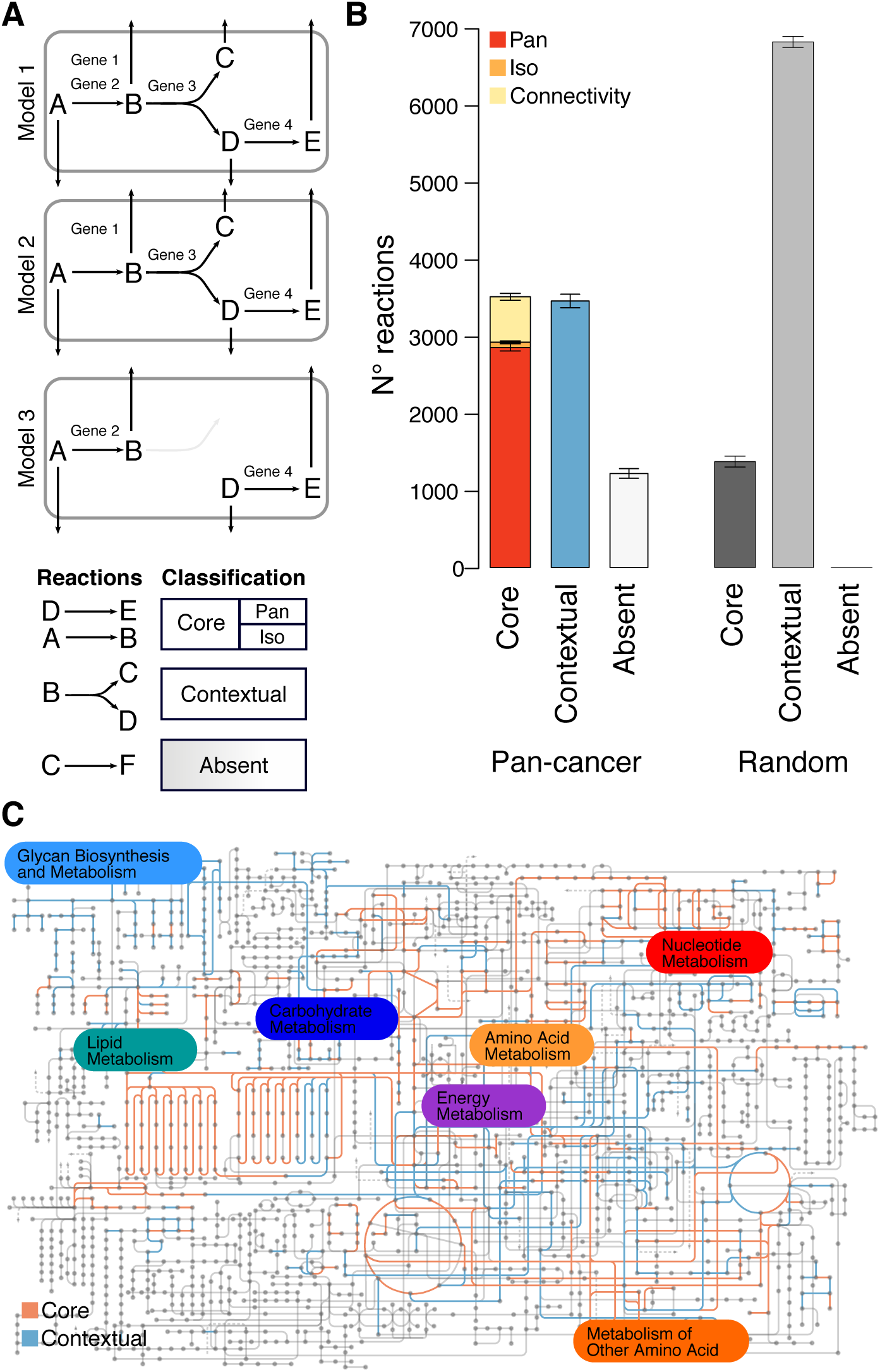
Classification of reactions based on their presence across 917 cancer genome-scale metabolic models. (A) Depiction of reaction classification. Core reactions are present in virtually all models, either by means of the same gene-reaction association (pan) or via different isoenzymes (iso). Contextual reactions are present only in a fraction of the models. Absent reactions are absent in virtually all models. (B) Number of core, contextual and absent reactions in the pan-cancer set and in a random set of 917 metabolic networks. The error bar represents the 95% confidence interval for the bootstrap statistics. See also Fig. S5. (C) KEGG map of human metabolism colored to distinguish core (red) from contextual (blue) reactions.

### Analysis of core metabolic reactions in cancer

Core reactions are expected to represent the backbone of cell metabolism. Thus, we compared core reactions as defined here to the list of reactions carried by the housekeeping proteome in normal human tissues based on Human Protein Atlas (HPA) data (*27*) (Fig. 3A). Excluding conn reactions, 2,900 out of 2,919 core reactions (99.3%) are housekeeping also in normal tissues, as expected. Noteworthy, cancer metabolism seemed to be generally associated with a large loss of function in metabolism, i.e. there were 2,083 housekeeping reactions in normal cells that were not present in any of the cancers. Furthermore, 19 reactions were core only in cancers. These were lumped in 5 reaction clusters, defined as sets of reactions encoded by the same gene(s): the mitochondrial conversion of arginine to ornithine and urea (encoded by *ARG2*); amino-acid transport (*SLC6A9* and *SLC6A14*); prostaglandin biosynthesis (*PTGS1* and *PTGS2*); sulfate transport (*SLC26A1, SLC26A2, SLC26A3, SLC26A7, SLC26A8, SLC26A9*); and ammonia transport (*RHAG*). Except for ammonia transport by *RHAG*, all other reactions have at least one associated gene that is highly expressed in the sample from which the model was reconstructed (Fig. 3B). At the same time, the corresponding proteins showed weak to no gene expression evidence in at least 30 out of 32 normal human tissues according to HPA (Fig. 3C). This is suggestive of a small acquisition in metabolic housekeeping functions in cancer compared to normal human cells, which should be otherwise considered entirely overlapping.

**Figure 3.**
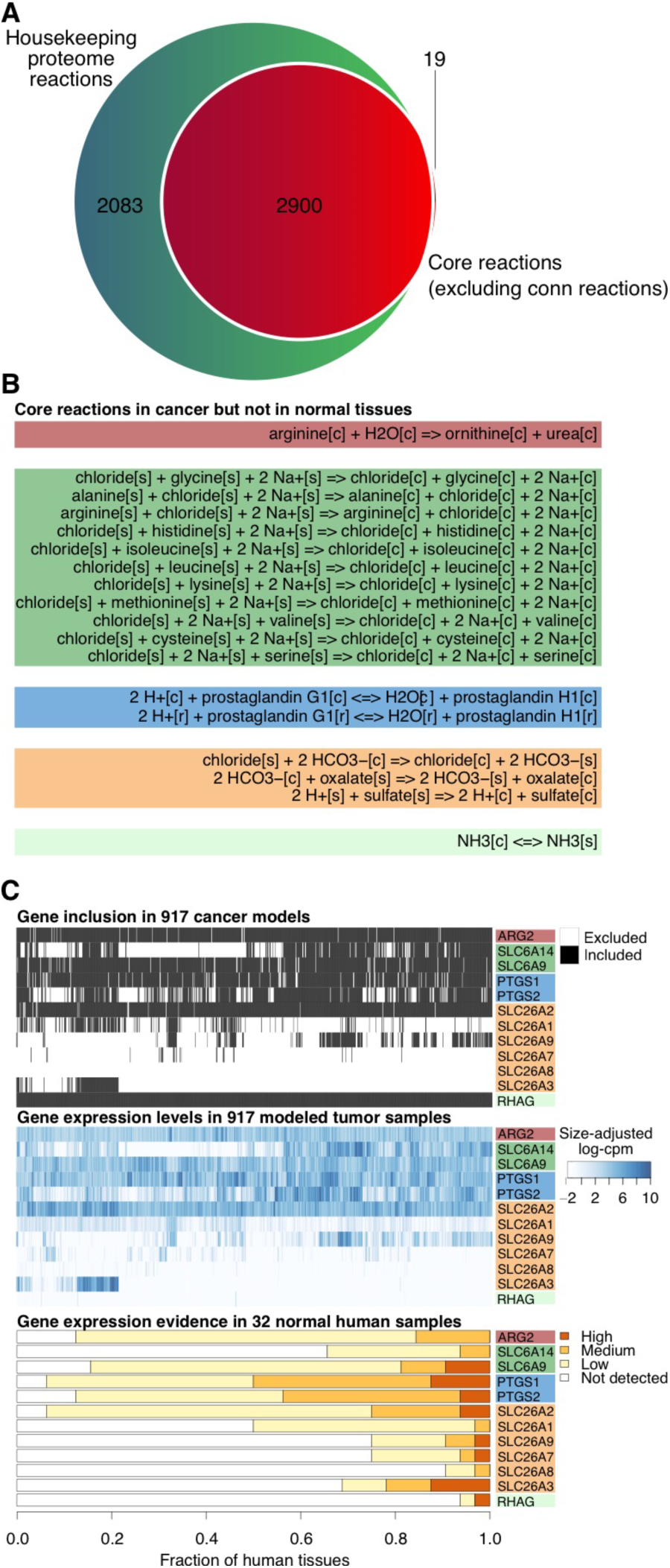
Core reactions in cancer versus reactions carried out by the housekeeping proteome in normal human tissues. A) Venn diagram of the reactions classified as core in this study (excluding conn-reactions) (red circle) or housekeeping on the basis of the gene expression pattern in 32 normal human tissues as reported in HPA (green circle). B) Core reactions in cancer but not housekeeping in normal tissues based on HPA data. C) Inclusion of core reactions not classified as housekeeping based on HPA data across the 917 cancer models and matched gene expression in the corresponding tumor samples. For comparison, the gene expression evidence in 32 normal human tissues for the genes encoding these reactions were retrieved from HPA.

Among the core reactions, we identified 69 core-iso reactions, which were included in all models but due to different gene-reaction associations. These were lumped in 16 reaction clusters (Fig. 4A). For example, the conversion of N-acetylputrescine to N4-acetylaminobutanal in the metabolism of arginine can be carried out in principle by 5 different amine oxidases (encoded by *AOC1, AOC2, AOC3, MAOA, MAOB*), whose expression varied considerably among samples for which GEMs had been reconstructed (Fig. 4B). We sought to identify if inclusion of a given isoenzyme in the GEM correlated with particular features of the modeled sample. In particular, we tested if the inclusion had a significant statistical association with the sample cancer type and on the presence of mutations in key cancer genes. We observed that in all cases the expression of a particular isoenzyme was affected by the sample cancer type, but in no instance by the presence of a particular mutated gene (likelihood ratio test, FDR < 0.001). This was well illustrated by the example of N-acetylputrescine degradation (Fig. 4C). This reaction is likely to be carried out by *AOC1*-*MAOA* gene products in rectal adenocarcinoma but *AOC3*-*MAOB* in breast invasive carcinoma. To provide a simplified yet exhaustive view of these associations, we computed for each cancer type the isoenzyme with the most frequent association with a given reaction cluster (Fig. 4A). Collectively, this result is consistent with the previously observed notion that some degree of diversity in gene inclusion across models was not reflected at the reaction level, due to the expression of cancer type-specific isoenzymes.

**Figure 4.**
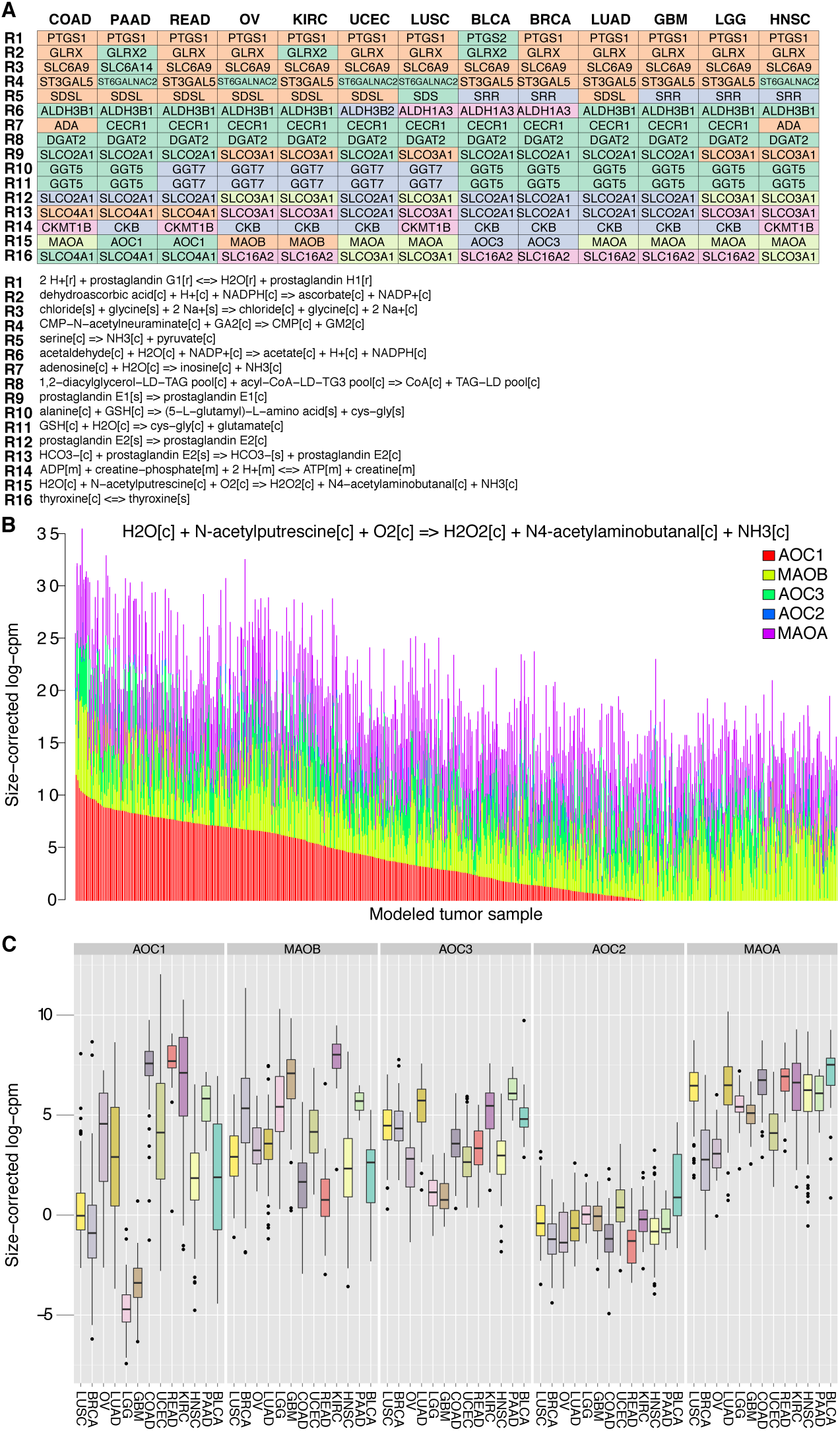
Core-iso reactions in cancer. Sixteen core-iso reaction clusters (i.e., set of reactions encoded by the same gene(s)) were identified following differential inclusion of isoenzymes in the different models. In all cases, differential inclusion was significantly associated with the sample cancer type. A) Most recurrent gene-reaction association in the models representing a given cancer type for the 16 core-iso reaction clusters (only one representative reaction shown per cluster). B) Expression plot for an example core-iso reaction, the conversion of N-acetylputrescine to N4-acetylaminobutanal in arginine metabolism, which is associated with 5 different genes (*AOC1*, *AOC2*, *AOC3*, *MAOA*, *MAOB*). Each bar stacks the expression levels of the 5 genes in the tumor sample from which the model was reconstructed. The bar of genes with size-adjusted log-cpm < 0 was neglected. Models were sorted according to *AOC1* expression. C) Expression boxplots for the 5 genes encoding the example reaction in B) when binned by cancer type. Keys as in Fig. 1.

### Analysis of contextual metabolic reactions in cancer

Contextual reactions are included only in a fraction of models. This can be explained in view of how the algorithm treated the input data. A reaction was prone to be excluded from a GEM in two cases: first, the expected expression of the associated gene(s) in the corresponding sample was not distinguishable from noise; or, the expected expression of genes associated with neighboring reactions in the corresponding sample was not distinguishable from noise, hence prejudicing the connectivity of the network. An example of contextual reactions is the ALOX12-mediated peroxidation of 12(*S*)-hydroperoxy-5*Z*,8*Z*,10*E*,14*Z*-eicosatetraenoic acid (12(S)-HPETE) to 12(S)-hydroxyeicosateraenoic acid (12(S)-HETE), a reaction in the metabolism of arachidonic acid that was included only in 163 out of 917 GEMs (17.8%). This was reflected by the expression level of *ALOX12* in the corresponding samples (Fig. 5A). Again, we sought to identify if inclusion of contextual reactions depended on the cancer type or on the presence of mutated genes in the sample. Out of 3,455, 3,269 reactions (94.6%) showed a significant association with the cancer type, while 1 reaction had a significant association with a mutated gene (likelihood ratio test, FDR < 0.001). As an example for the former case, the cancer type-dependency of the ALOX12-catalyzed reaction was evident from *ALOX12* expression level in the different types, which was particularly high in head and neck and lung squamous cell carcinomas and in bladder adenocarcinomas (Fig. 5B). As for the latter case, the only mutated gene-associated reaction is the mitochondrial hydrolysis of glutamine to glutamate encoded by glutaminase-2 (*GLS2*), which was excluded preferentially in models for tumors in which *TP53* was mutated (Fig. S7).

**Figure 5.**
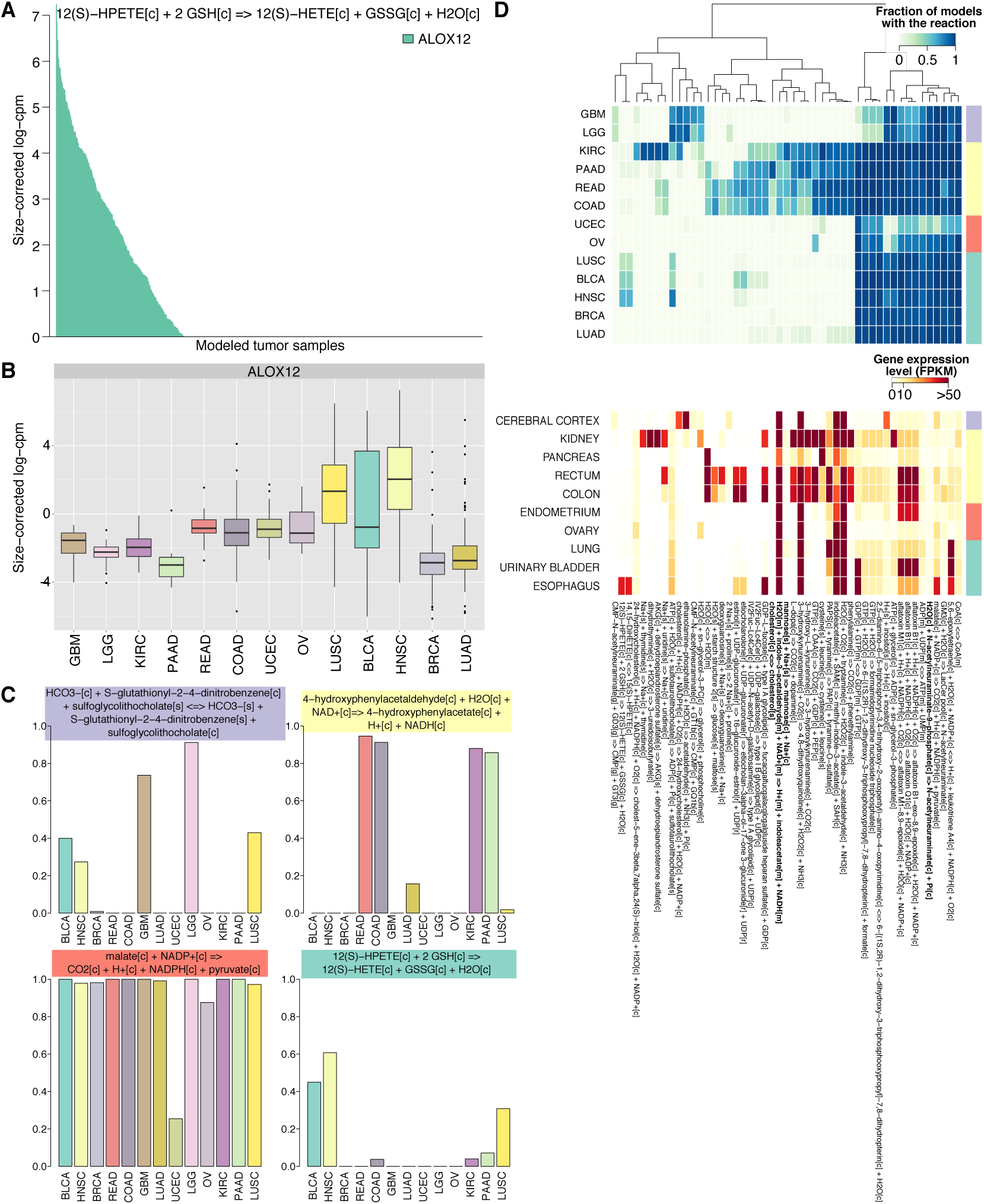
Contextual reactions in cancer were mostly dependent of the cancer type and some of these may represent specific gain or loss of metabolic functions compared to the matched normal tissues. A) An example of contextual reaction, the conversion of 12-HPETE to 12-HETE by arachidonate 12-lipoxygenase, ALOX12), and its expression plot across models. B) Expression boxplots for *ALOX12* binned by cancer type. C) Barplots for the fraction of models from a cancer type (columns) that contain 4 representative contextual reactions (rows) for each cancer-type consensus clusters (horizontal colored bars). D) Heatmap of the fraction of models from a cancer type (columns) that contained the 49 most representative contextual reactions (rows) for the four cancer-type consensus clusters (see Fig. S8, colored horizontal bars). A blue entry indicates that all models belonging to that cancer type included a given reaction (white if vice versa). Below, maximum expression detected in matched normal tissues among the genes encoding for each representative contextual reaction. Keys as in Fig. 1.

In order to reduce the complexity of cancer-type dependency in contextual reactions, we computed for each cancer type the fraction of GEMs where the reaction was included (e.g. 0 means that no GEMs from samples belonging to a given cancer type included that contextual reaction). Next we performed consensus hierarchical clustering to detect consensus clusters of cancer types where contextual reactions appeared to segregate (Fig. S8). Four clusters were identified: brain tumors (low grade glioma and glioblastoma multiforme); kidney-pancreatic-colorectal tumors (clear cell renal cell carcinoma and pancreatic, colon, and rectum adenocarcinoma); female reproductive system tumors (ovarian cancer and uterine corpus endometrial cancer); other tumors (bladder and lung adenocarcinomas, breast carcinoma, head and neck and lung squamous cell carcinomas). Finally, we performed random forest cross-validated variable selection to select contextual reactions that exhibited the strongest representativeness to either cluster. We found 49 representative contextual reactions. For example, the transport of sulfoglycolithocholate was preferentially included in brain tumor GEMs (mean in cluster, 82.4%; outside cluster, 19.7%); the conversion of phenylalanine to phenethylamine was almost exclusively present in kidney-pancreatic-colorectal tumor GEMs (in cluster, 89.9%; outside, 1.9%); the conversion of cytosolic malate to pyruvate was less common in female reproductive system tumor GEMs (in cluster, 56.4%, outside, 99.3%); and the above-seen conversion of 12(S)-HPETE to 12(S)-HETE was mostly included in other tumor GEMs (in cluster, 27.3%, outside, 1.9%) (Fig. 5C).

Since virtually all contextual reactions displayed cancer-type dependency, which in turn appeared to cluster by human tissue similarity, we interrogated whether the context-specificity of these 49 representative reactions simply mirrored the expression patterns of the associated genes in the normal tissues of origin, as previously suggested for metabolic genes (*3*, *5*). Therefore, we retrieved RNA-seq gene expression data from HPA for all putative tissues of origin for the cancer types in this study (except breast, unavailable). We then extracted which gene had the maximum expression level in each tissue among all genes associated with each reaction. As hypothesized, the expression patterns in normal tissue closely resembled the availability of a given reaction in the GEMs of their malignant counterpart (Fig. 5D). However, we noticed some exceptions, both in terms of gain and loss of metabolic function. As an example of gains of function, the conversion of N-acetylneuraminate-9-phosphate to N-acetylneuraminate was present in all GEMs except in some belonging to the cluster of female reproductive system tumors. However, the associated gene, *NANP*, is not expressed in any major normal tissue (range, 0 to 4 FPKM). As a second example, the sodium-dependent transport of mannose and the absorption of cholesterol were both featured in GEMs belonging to the cluster of kidney-pancreatic-colorectal tumors. However, the encoding genes, *SLC5A9* and *NPC1L1* respectively, are not expressed in the matched normal tissues, but only in the duodenum and small intestine (range, 52 to 55 FPKM and 41 to 45 FPKM resp.). An example of loss of function was the mitochondrial oxidation of indole-3-acetaldehyde to indoleacetate in tryptophan catabolism, which is catalyzed by several NAD-dependent aldehyde dehydrogenases with wide specificity (ALDH1B1, ALDH2, ALDH3A2, ALDH7A1, ALDH9A1). While these enzymes are largely expressed in all major normal tissues, the catalyzed reaction was mostly absent in all cancer GEMs, except for a fraction of models belonging to the cluster of kidney-pancreatic-colorectal tumors.

In conclusion, contextual reactions represented on average 26% of all metabolic reactions in any given cancer network. This analysis revealed that the context-specificity is often related to the cancer type. In particular, most contextual reactions segregated in four tissue clusters and, with few notable exceptions, this pattern resembled the expression profiles of the cancer tissue of origin.

### Expression of contextual reactions correctly distinguished cancer type clusters

Next we sought to validate the 49 contextual reactions here found to distinguish the metabolic networks of four cancer type clusters defined above. To this end, we retrieved expression data from TCGA for 66 genes encoding these contextual reactions in an independent cohort of 4,462 tumor samples spanning the same 13 cancer type. Next, we trained a random forest classifier on 2,231 samples (*28*) to assign samples to one of the four cancer type clusters based on the expression level of this 66 gene signature (Fig. S9A). We then tested the so-trained classifier on the remaining 2,231 samples (Fig. S9B). The out-of-bag error was 3.38% in the training set and 2.9% in the test set. The multiclass area-under-the-curve (AUC), as defined by Hand and Till (*29*), was 0.973.

The classification performance might be biased due to inherent differences in the gene expression profile of the tissue of origins in the four clusters. In other words, any set of 66 genes could distinguish the four clusters because their expression is supposedly modulated in a tissue-specific fashion in healthy conditions. To control for this, we repeated the above multiclass classification using 1,000 random 66 gene-signatures and verified that the classifier based on the expression of contextual reactions outperformed the random classifier in 98.8% cases (permutation test *p* = 0.012, Fig. S9C). This suggestive that not to any set of 66 genes is differentially regulated between these cancer type clusters simply because of inherent differences in expression in their respective tissue of origins. To further check whether these 66 genes are differentially expressed between cancer type clusters, but yet substantially similar in their corresponding normal tissues, we performed differential expression analysis using a generalized linear model to fit gene expression to the cancer type cluster while controlling for the matched-normal tissues of origin. We retrieved 438 tumor-adjacent normal samples from TCGA for the same cancer types as above except brain tumors, for which only 5 normal samples were available (sample size range per cluster: 24 - 287). Thus, we analyzed gene expression for the three remaining cancer type clusters (accounting for 3,867 tumor samples, sample size range per cluster: 459-2452). We observed that 64 of 66 genes in the signature (97%) displayed differential regulation between cancer type clusters (FDR < 0.001). For example, *AOC1* is moderately expressed in the female reproductive system tumor cluster, but lowly expressed in the other tumor cluster (FDR = 9*10^−69^, Fig. S9D). For 48 of 66 genes, the expression was also significantly different from at least one cluster of matched normal samples. However, importantly, we found no evidence of expression difference between the corresponding clusters of matched normal samples (FDR > 0.01). For example, *AOC1* expression is overexpressed in female reproductive system tumors compared to matched normal samples (FDR = 2*10^−4^). But normal samples from the female reproductive system had similar *AOC1* expression to normal samples from tissues matched to the other tumor cluster (FDR = 0.988). Besides *AOC1*, we computed the most significant gene for the remaining two cluster comparisons: *ALDH2*, repressed in the other tumor cluster compared to kidney-pancreatic-colorectal tumor cluster, but similarly expressed in the respective tissue of origins; and *CYP3A5*, very lowly expressed in female reproductive system tumor cluster compared to the kidney-pancreatic-colorectal tumor cluster, yet again similarly expressed in the respective tissue of origins (Fig. S9D).

In conclusion, even though gene expression analysis is not a direct evidence of the occurrence of the underlying reactions and fails to account for the systems of metabolic reactions in which a gene is expressed, these results in a large independent cohort suggest that the 49 contextual reactions in the distinct cancer type clusters arise from cancer type-specific regulation of the corresponding genes that are not attributable to substantial differences in the gene expression profiles of the matched tissue of origins, arguing that they may represent cancer type-specific metabolic liabilities.

## DISCUSSION

Reprogramming of cellular metabolism is a hallmark of malignant transformation, as suggested by accumulating evidence that uncovered tumor-specific molecular mechanisms of metabolic regulation (*1*, *2*). However, only few studies explored tumor metabolism at the systems level (*7*). Since networks are the natural structure of complex systems like metabolism, in this study we adopted a network-centric approach to characterize the landscape of metabolic reactions expressed in different cancers. Despite the substantial diversity of the genes included in each model across cancers, the resulting metabolic networks were strikingly similar in terms of included reactions, probing the robustness and redundancy of the human metabolic network.

The overwhelming majority of core reactions in this study are carried out by enzymes classified as housekeeping in normal tissues. This argues that cancer cells, regardless of their origin, vastly maintain the backbone of cellular metabolism and no specific metabolic function is universally acquired as a result of the transformation. Potential exceptions to this could be represented by the residual core reactions, which are so-classified only in cancer metabolism networks. While inclusion of these reactions in all GEMs certainly depends on network connectivity and functionality, we also observed an obvious correlation with gene expression in the underlying cancer samples. There could be many alternative biological reasons why these reactions are universally present in cancer metabolic networks while being seldom expressed in normal human tissues. Perhaps an intriguing pattern is the role of both *ARG2* and *PTGS1/2* in the generation of metabolites that control endothelial cell proliferation (*30*, *31*) and activation of the immune system (*31*, *32*), which are likely to be dispensable metabolic functions in the majority of healthy human cells. Another interesting observation is the presence of core-iso reactions, included in all models because at least one isoenzyme was expressed in the modeled tumor sample. Core-iso reactions were previously observed by Hu *et al.*, for example in the case of aldolase isoenzymes (*ALDOA, ALDOB*, *ALDOC*) (*3*). Here, we complemented these findings and verified that in all cases the differential expression of isoenzymes could be attributed to differences in cancer type between the modelled samples. For example, oxidation of primary amines (like acetylputrescine) is possible in all models because at least one among the spectrum of monoamine oxidases (*MAOA* and *MAOB*) and amine oxidases (*AOC1, AOC2, AOC3*) was expressed in each underlying sample.

Contextual reactions were present only in a fraction of the reconstructed metabolic networks. In one case, which is the mitochondrial synthesis of glutamate from glutamine encoded by *GLS2*, the contextual reaction was strongly associated with absence of *TP53* mutations in the modeled sample (regardless of its cancer type). This result is strikingly concordant with the notion that p53 directly controls *GLS2* expression in either stressed or not stressed conditions (*33*) and further suggests that the presence of the underlying reaction is partly dictated by mutations in *TP53* independent of the cancer type. In 94.6% of the other cases, the inclusion of contextual reactions could be attributed to differences in cancer types, which we narrowed down to differences across four cancer type clusters. In particular, 49 contextual reactions were strongly representative for the four cancer type clusters because of differential regulation of the associated genes across cancers of different types, irrespective of the expression level in the corresponding tissues of origin. At the same time, in general, the pattern of contextual reaction inclusion in a model largely resembled the expression pattern of the associated gene(s) in the matched normal tissue of origins. Exceptions to this were noted for few reactions, for example the mitochondrial oxidation of indole-3-acetaldehyde to indoleacetate in tryptophan catabolism, by several NAD-dependent aldehyde dehydrogenases (ALDH1B1, ALDH2, ALDH3A2, ALDH7A1, ALDH9A1). This case is intriguing because of *ALDH2*: its cancer type-cluster specificity and the substantial expression difference with the matched normal tissues seem both to stem from the underlying patterns of *ALDH2* expression (Fig. S9D). *ALDH2* is moderately to highly expressed in most normal samples, but it is strongly repressed in all cancers except those belonging to the kidney-pancreatic-colorectal cluster, as observed in our network-centric approach (Fig. S9C). For the other cases, we could not derive equally straightforward relations with gene expression, likely due to the complexity of metabolic network around these reactions. An interesting outcome of this analysis is that the expression of the 66 genes encoding these 49 contextual reactions could accurately classify tumor samples into four groups of cancer types. This clearly points to metabolic similarities between certain cancer types, which can be used for identification of common drug targets for different tumor types that cluster in this analysis.

A surprising result was the apparent enrichment of neurological metabolic functions in cancer compared to normal samples. This was evident at different points: the core status of transport reactions catalyzed by solute carrier 6 gene family (the above mentioned *SLC6A9* and *SLC6A16*), which are traditionally classified as neurotransmitter transporters (*34*) and indeed were not found to be housekeeping in normal tissues; the differential expression of monoamine oxidases (*MAOA* and *MAOB*) in different cancer types possibly to guarantee primary amine oxidation as a core-iso reaction, even though these oxidases are classically linked to the metabolism of neuroactive amines; and the inclusion the *NANP*-catalyzed reaction in all models except in some of those in the female reproductive system tumor cluster, which generates N-acetylneuraminate (or Neu5Ac), a critical component of neuronal membranes. These observations are at odds with the notion that tumors lack innervation, but other explanations for their expression in the metabolic network are possible. On one hand, all these enzymes might fulfill other less characterized metabolic functions. On the other hand, it has already been observed aberrant expression of neuronal genes in non-brain tumors, which has been attributed to not well understood survival advantages (*35*-*37*), but define, for example, an established phenotype of prostate cancers (*38*).

Modeling metabolic networks at genome scale comes with challenges, mostly arising from knowledge gaps in the biochemistry of several metabolic reactions (*39*). The gene-reaction associations in this study were occasionally inferred, since the specificity and *in vivo* activity of some enzymes is unknown. Even though we drew our conclusions based on stringent statistical models, these shortcomings may introduce a systematic bias that is difficult to estimate and control for in downstream analysis. Hence, we expect that some quantitative claims will be revisited in light of improved knowledge on the biochemistry of human metabolism. An alternative approach could be (differential) expression analysis for each gene associated with a metabolic reaction, as we and others have implemented previously (*3*-*5*). However, we adopted a network-centric approach because reactions might be expressed or not depending on the availability of the neighboring reactions or, in a broader context, of a metabolic pathway or even a metabolic function; or they could be carried out by differentially expressed isoenzymes. This context is lost in canonical gene expression analyses. The network structure may occasionally override information coming from gene expression data. This provides an opportunity for discovery, but at the same time complicates the interpretability of certain results in light of gene expression evidence, as in the case of *RHAG* expression in Fig. 3C. This can also result in the opposite situation, i.e. metabolic genes seemingly expressed in the tumor yet absent at the reaction level because either unconnected or supporting an unknown metabolic function. Another challenge is the paucity of large-scale cell type-specific data. Despite the TCGA requirement that a sequenced tissue should contain >80% of tumor nuclei, its reconstructed network does not necessary reflect the reactions occurring in a single cancer cell nor it can dissect their distribution in the different clones in the tumor. Rather it compiles all the reactions expressed in the cell population of that tumor. Whilst a limitation, this acknowledges the contribution of stromal and immune cells to the metabolic plasticity of tumor (*40*).

In conclusion, our findings suggest that tumors express a metabolic network of core reactions with housekeeping functions and cancer type-specific contextual reactions that are virtually overlapping with the corresponding normal tissue of origin. In light of this vast similarity, metabolic reprogramming implicated with cancer transformation might just reflect the plasticity of tumors to adapt to varying environmental and genetic factors by leveraging on the complexity of the metabolic network. This also suggests that targeting tumor metabolism may result in either toxicity or resistance, because the underlying metabolic network supports essentially the same metabolic functions as in the matched normal tissue. Nonetheless, exceptions to this similarity with normal tissues were uncovered in this study, and we believe that this is a valuable resource to further investigate selective liabilities of tumor metabolic networks.

## MATERIALS AND METHODS

### Reconstruction of tumor genome-scale metabolic models

Read count tables for 18,956 genes from 1,082 primary tumor samples were retrieved from The Cancer Genome Atlas (TCGA), for which both gene expression and mutation data were simultaneously present: These samples were sequenced using Illumina HiSeq or Genome Analyzer RNA-seq platforms. For each sample, counts were transformed into log-counts-per-million (log-cpm) and multiplied with a normalization factor that accounts for differences in library sizes across samples, which was calculated using the *edgeR* R-package (*41*). The resulting size-adjusted log-cpm for a gene is an expression level measurement comparable across samples (*42*). The probability that a gene is truly expressed in an individual sample was estimated using a Bayesian statistical framework by computing the probability that its expected distribution of read counts in that sample is better explained by the expected distribution of read counts of genes that should not be expressed in that samples. The expected size-adjusted log-cpm for gene *i* in sample *j* was calculated based on specific features of sample *j* (like the belonging to a certain cancer type or the presence of a mutation in a key gene) using a pre-specified generalized linear model (*1*) from(*43*):

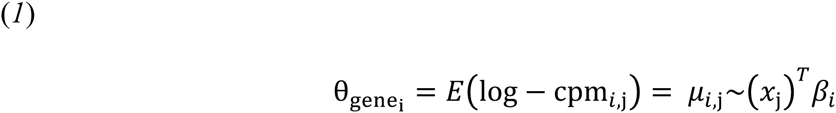

where the expected size-adjusted log-cpm for gene *i* in sample *j* is equal to the linear combination of sample feature variables (*x*_j_) and coefficients for the effect of that feature on gene *i* (β_*i*_). The coefficients were fitted for each gene to the observed size-adjusted log-cpm across the 1,082 samples by ordinary least square regression using *voom* R-package (*42*). The linear model for the expected expression of gene *i* is referred to as θ_gene_i__. We reasoned that gene *i* is truly expressed in sample *j* if its observed size-adjusted log-cpm is better explained by a normal distribution with parameters derived from θ_gene_i__, rather than by linear models constructed on the expected size-adjusted log-cpm of genes that are not supposed to be expressed in sample *j*. We selected 7 testis-specific gene products as “noise” genes: *ACRV1, ADAM2, BOLL, DKKL1, FMR1NB, TEX101*, and *ZPBP2*. These proteins are not detected in any other tissues according to the Human Protein Atlas (*27*), yet reads that align to their loci are seldom detected by the RNA-seq platform. The posterior probability that gene *i* is truly expressed in sample *j*, or in other words the likelihood of the linear model θ_gene_i__. for gene *i* being better than alternative linear models θ_gene_g__., was than calculated using the Bayesian formula (*2*):

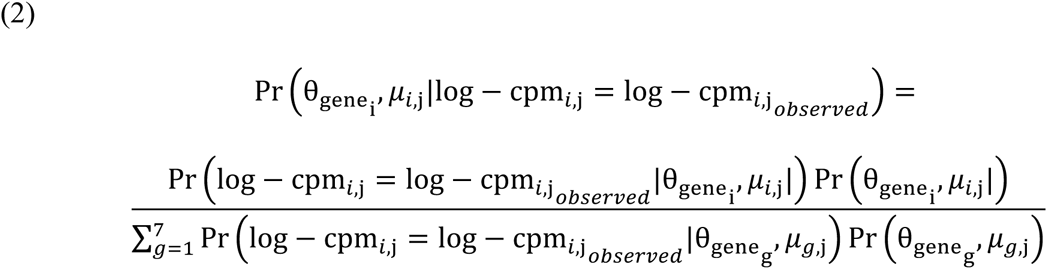

where the prior probability of the linear model θ_gene_i__ was set equal to 50% and the prior probabilities for each alternative linear model θ_gene_g__ derived from each of 7 “noise” genes was equally split so that they would sum to 50%. The output is a *m* × *n* matrix of posterior probabilities *P*, where *m* is 18,956 genes and *n* is 1,082 samples, and *P*_*m,n*_ ∈ (0,1).

The posterior probabilities’ vector for sample *j* was used to score each reaction in the automatic reconstruction of a genome-scale metabolic model (GEM) from the reference generic human GEM, *HMR2. HMR2* contains 8,184 reactions associated with 3,765 genes. The reconstruction was performed using the tINIT algorithm, as described previously (*24*). Briefly, the algorithm implements a mixed-integer linear problem (MILP) that, starting from a reference metabolic network (in this case *HMR2*), maximizes the inclusion in the final model of reactions that are positively scored while maximizing the exclusion of reactions that are negatively scored. Other constraints in the MILP guarantee that the final model features a connected and functional network, in that it can simulate a flux in each reaction under standard medium conditions, and it can simulate a flux through 56 fundamental metabolic tasks, including biomass growth (*24*). In each reconstructed sample *j*, the reaction score was equal to the posterior probability of the associated gene in sample *j*, as returned from (*2*). If more than one gene was associated with a reaction, the most positive score was retained because indicative that the reaction can be carried out by at least one expressed gene product, as previously implemented (*25*, *44*). We selected a threshold equal to 0.99 that we subtracted to the posterior probabilities’ vector for sample *j*, before scoring the reactions, so that only genes with a posterior probability > 99% to be truly expressed (as defined above) have positive values and will tend to be included by tINIT. We selected this threshold as an arbitrary metric reflecting 99% statistical confidence, which we reasoned to be a conceptually superior over arbitrary thresholds based on expression scores as previously implemented in automatic reconstruction of GEMs (*25*, *45*, *46*). The parameters for the reconstruction were identical for all samples, in particular the maximum allowed relative gap in the optimal solution returned from the MILP was set to 5%. The reconstruction was implemented in Matlab 7.11 using Mosek v7 as solver.

### Quality control of reconstructed tumor GEMs

We evaluated the quality of the reconstruction according to the following criteria: convergence to an optimal solution; the accuracy of gene inclusion should be > 90%; the corresponding Matthews correlation coefficient, an unbiased estimator of accuracy, should be > 0.80; and the number of reactions added during the reconstruction in order to be able to perform the 56 metabolic tasks considered in the tINIT algorithm should be < 1% of all reactions included in the model. These thresholds were selected to prevent reconstructions of questionable quality to be included in downstream analysis, in consideration that most reconstructions converged successfully and only some outliers fall below the selected values (Figure S1). The accuracy for model *k* derived from sample *j* was calculated in terms of true/false positives/negatives in the final model *k*, where a true positive is a gene with posterior probability > 99% in sample *j* and included in model *k* and a true negative is a gene with posterior probability <= 99% in sample *j* and excluded in model (false positive/negative if vice versa). Of 1,082 initially considered samples, 917 GEMs could be reconstructed that satisfied the above criteria. Principal component analysis (PCA) to evaluate similarity of the 917 reconstructed GEMs was performed on the gene/reaction inclusion matrix, where each row is a gene/reaction in the reference GEM, *HMR2*, and each column represents a reconstructed GEMs. A matrix cell is equal to 1 if the corresponding *HMR2* gene/reaction is included in the corresponding reconstructed GEM. PCA was performed on unscaled data with mean centering using the *ade4* R-package (*47*).

### Reaction classification and analysis

A *HMR2* reaction was classified as “core” if included in > 95% of all reconstructed GEMs; “absent” if included in less < 5% of all reconstructed GEMs; “contextual” if otherwise. We selected these thresholds because the underlying distribution seemed to reach a plateau at these values (Fig. S5). Core reactions were further sub-classified into “pan” if their inclusion was due to inclusion of the same associated gene(s) in > 95% of all reconstructed GEMs and “iso” if otherwise, meaning that different isoenzymes are present in different models for that core reaction. We estimated how robust was the number of core, contextual and absent reactions to potential outliers in the set of reconstructed GEMs by bootstrapping the count of each category 1,000 times. We computed the count of core, contextual and absent reactions in 1,000 sets of 917 random GEMs, reconstructed by including *v* random genes from the gene-reaction association matrix of *HMR2*, where *v* is a 1x917 vector equal to the number of genes included in the actual set of 917 GEMs reconstructed using tINIT. Core and contextual reactions were mapped to the generic KEGG metabolic map using iPath2 (*48*) (if univocally identified using KEGG IDs) and grouped according to *HMR2* metabolic sub-systems.

Core reactions in tumors were compared to housekeeping reactions in normal tissues, so classified based on Human Protein Atlas (HPA) evidence for the associated genes. We compiled this list by including genes that were either expressed (> 0.5 FPKM) in all 32 normal tissues considered by HPA (“expressed in all” HPA category); or in > 30 normal tissues; or expressed in a subset of the 32 tissues that share little functional similarity, e.g. *CBLN3* in cerebellum, lung and small intestine, which suggests that the gene may be expressed promiscuously across human tissues (“mixed” HPA category). We counted 69 core-iso reactions, which could be lumped in 16 reaction clusters, sets of reactions encoded by the same gene(s). We tested whether the preferential inclusion of an isoenzyme in certain model could be attributed to the cancer type or presence of mutations in 9 key cancer-associated genes, which were featured in the linear model (*1*), using a likelihood ratio test. After adjustment for multiple testing (*49*), an association was considered significant if the false discovery rate (FDR) < 0.001. The most recurrent gene-reaction association present in models belonging to a certain cancer type was chosen as the most likely isoenzyme to carry out the core-iso reaction in that cancer type.

We tested if the preferential inclusion of a contextual reaction in a certain model could be attributed to the cancer type or presence of mutations in 9 key cancer-associated genes, using a likelihood ratio test. Since 94.6% of contextual reactions showed an association with cancer type, we sought to reduce the complexity by deriving clusters of cancer types where a contextual reaction is included more or less frequently than expected. We performed consensus clustering on a *l* × *c* matrix, where *c* is the number of cancer types (13) and *l* is the number of contextual reactions (3,269), and each matrix cell is the fraction of models derived from samples of the corresponding cancer type containing the corresponding contextual reaction. The optimal number of cluster *w* = 4 was selected based on the observation that for *w* > 4 the relative change in the area-under-CDF-curve was < 0.1, where CDF stands for the empirical cumulative distribution function of consensus distributions for up to *w* clusters. Consensus clustering was implemented using *ConsensusClusterPlus* R-package (*50*). The 49 contextual reactions most representative of the four clusters were chosen based on variable selection procedure applied to a random forest classifier constructed to assign a model to either of the four consensus clusters based on inclusion of contextual reactions. Random forest-driven variable selection was implemented using the *varSelRF* R-package (*51*).

### Validation of cancer type cluster-dependency of contextual reactions

Gene expression profiles for 4,462 primary tumor samples from the same 13 cancer types (sample size per type: 94 to 978) were retrieved from TCGA and transformed to size-adjusted log-cpm as described above. The profiles were randomly split in a training and test set so that 50% of samples belonging to a certain cancer type cluster were included in each set. A random forest classifier was constructed on the expression level in the training set of 66 genes, selected because associated with the 49 contextual reactions most representatives of the different clusters (see above). The performance in the classification of samples in the test set to the 4 cancer type clusters based on the 66 gene expression signature was evaluated by calculating the multiclass AUC (*29*). The same procedure was repeated for 1,000 random 66 gene expression signatures. The random forest classification was implemented using the *RandomForest* R-package (*28*). Differential gene expression analysis for the 66 gene expression signature was performed. Samples belonging to the brain tumor cluster because not enough tumor-adjacent normal samples were found to match these cancer types in TCGA. We retrieved 438 normal samples from TCGA that matched the remaining cancer types. We evaluated differential gene expression between tumor vs. normal samples belonging to the same cluster, between tumor samples belonging to different clusters, and between normal samples belonging to different clusters. After adjustment for multiple testing, a gene was considered differentially expressed between groups if FDR < 0.001. The differential gene expression analysis was implemented using the *voom* and *limma* R-packages (*42*, *52*)

## Acknowledgments

The authors wish to thank Leif Väremo for his contributions on the reaction classification, and Avlant Nilsson, Benjamín José Sánchez, and Michael Gossing for a critical review of the manuscript. The computations were performed on resources provided by the Swedish National Infrastructure for Computing (SNIC) at C3SE. Knut and Alice Wallenberg Foundation is acknowledged for financing this work.

## SUPPLEMENTARY FIGURES

**Figure S1.**
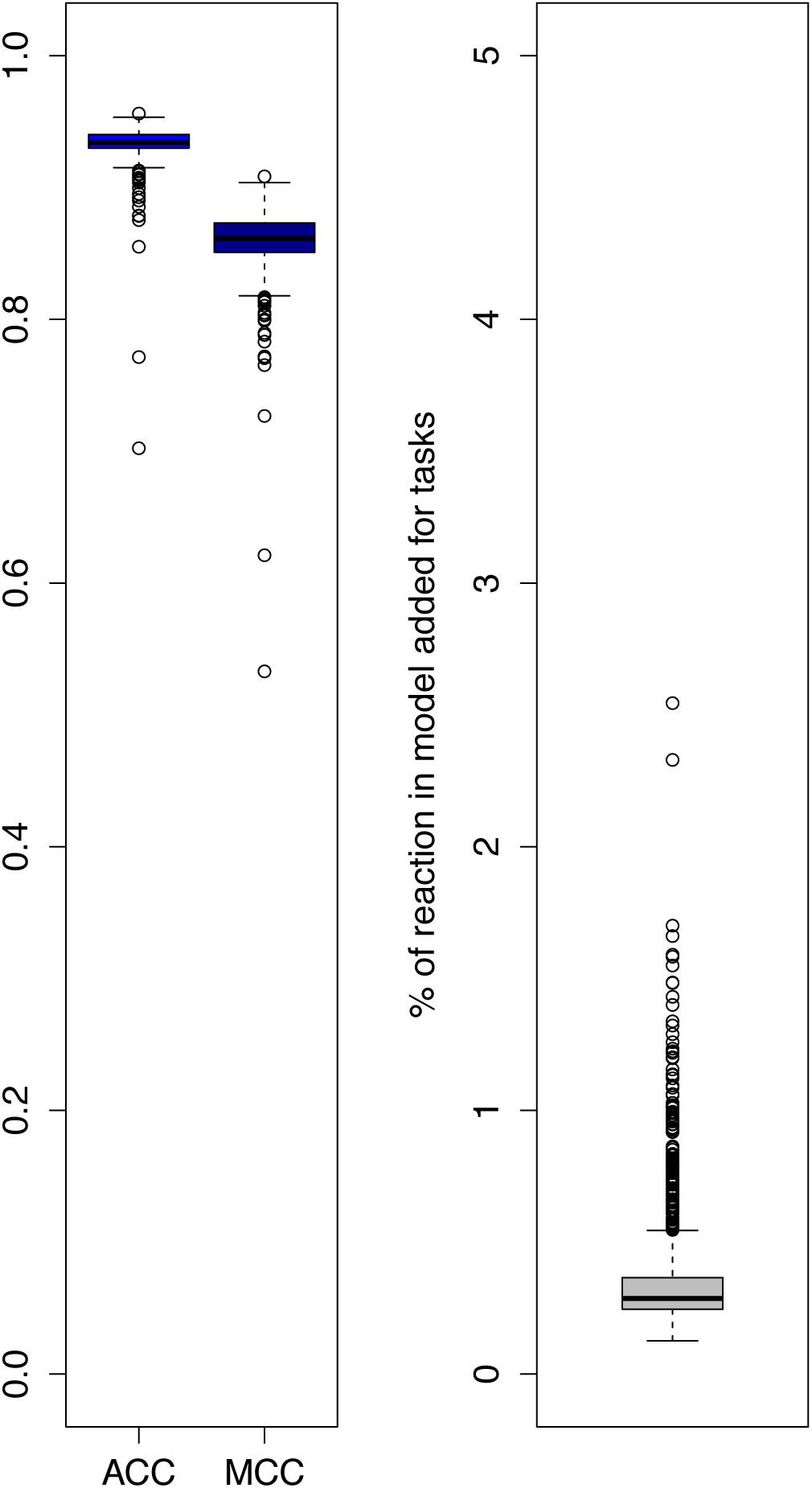
Accuracy of the model reconstructions in terms of fraction of the sum of genes that should be included and were included in the model (true positives) and genes that should be excluded and were excluded from the model (true negatives). Left: boxplots of accuracy (ACC) and Matthews Correlation Coefficients (MCC) for all model reconstructions. Right: percentage of reactions in the model enforced to fulfill a predefined list of metabolic tasks.

**Figure S2.**
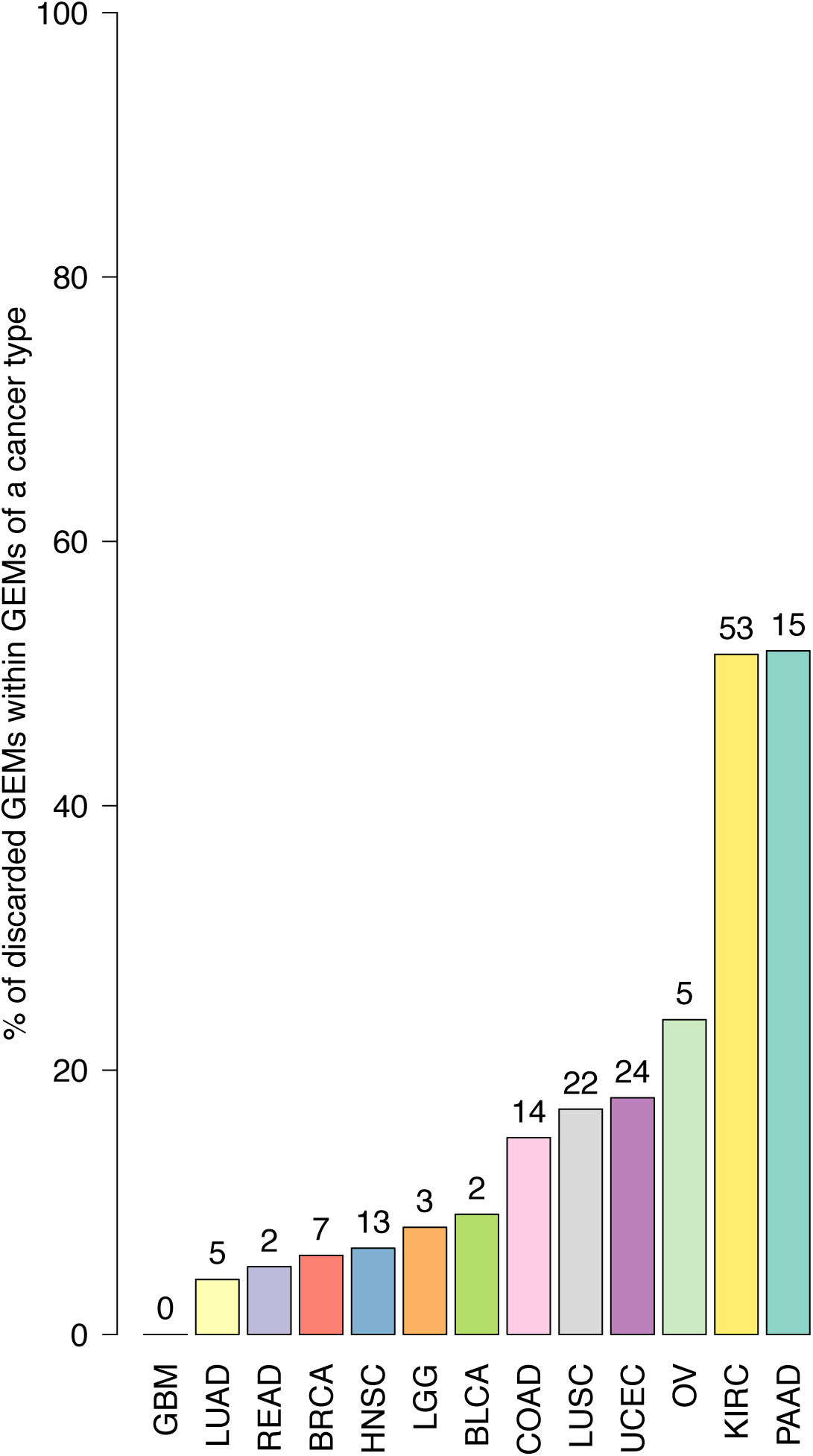
Percentage of models belonging to a given cancer type that did not meet the reconstruction quality criteria. Absolute number of discarded models is shown on top of each bar. Key: BLCA – Bladder adenocarcinoma, BRCA – Breast carcinoma, COAD – Colon adenocarcinoma, HNSC – Head and neck squamous cell carcinoma, GBM – Glioblastoma multiforme, KIRC – Clear cell renal cell carcinoma, LGG – Low grade glioma, LUAD – Lung adenocarcinoma, LUSC – Lung squamous cell carcinoma, OV – Ovarian carcinoma, PAAD – Pancreatic adenocarcinoma, READ - Rectum adenocarcinoma, UCEC – Uterine corpus endometrial carcinoma.

**Figure S3.**
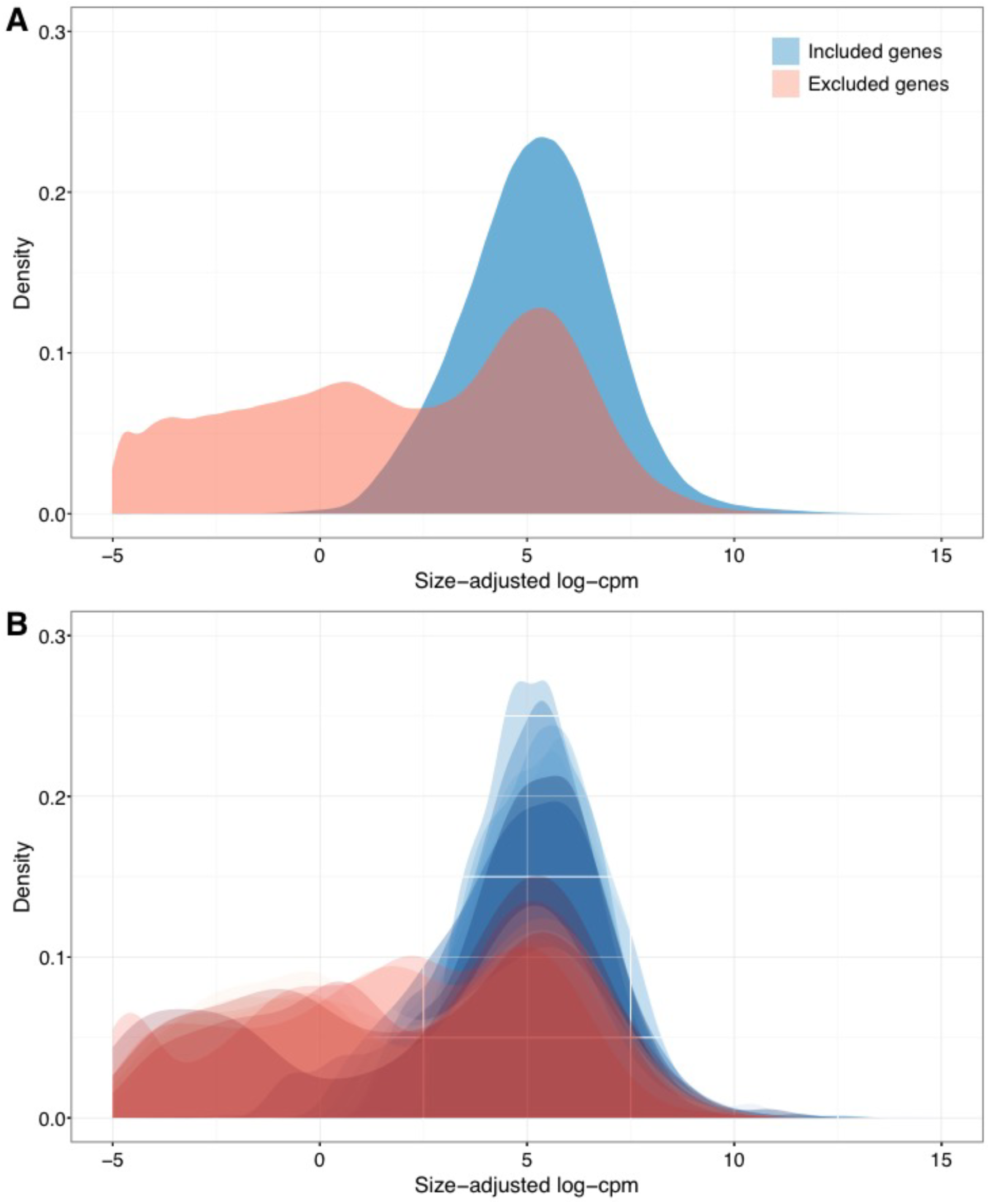
Density plots of gene expression values (in log_2_ counts-per-million, adjusted for the library size) for the genes excluded (red) or included (blue) in the reconstruction of all models (A) or in each model (B).

**Figure S4.**
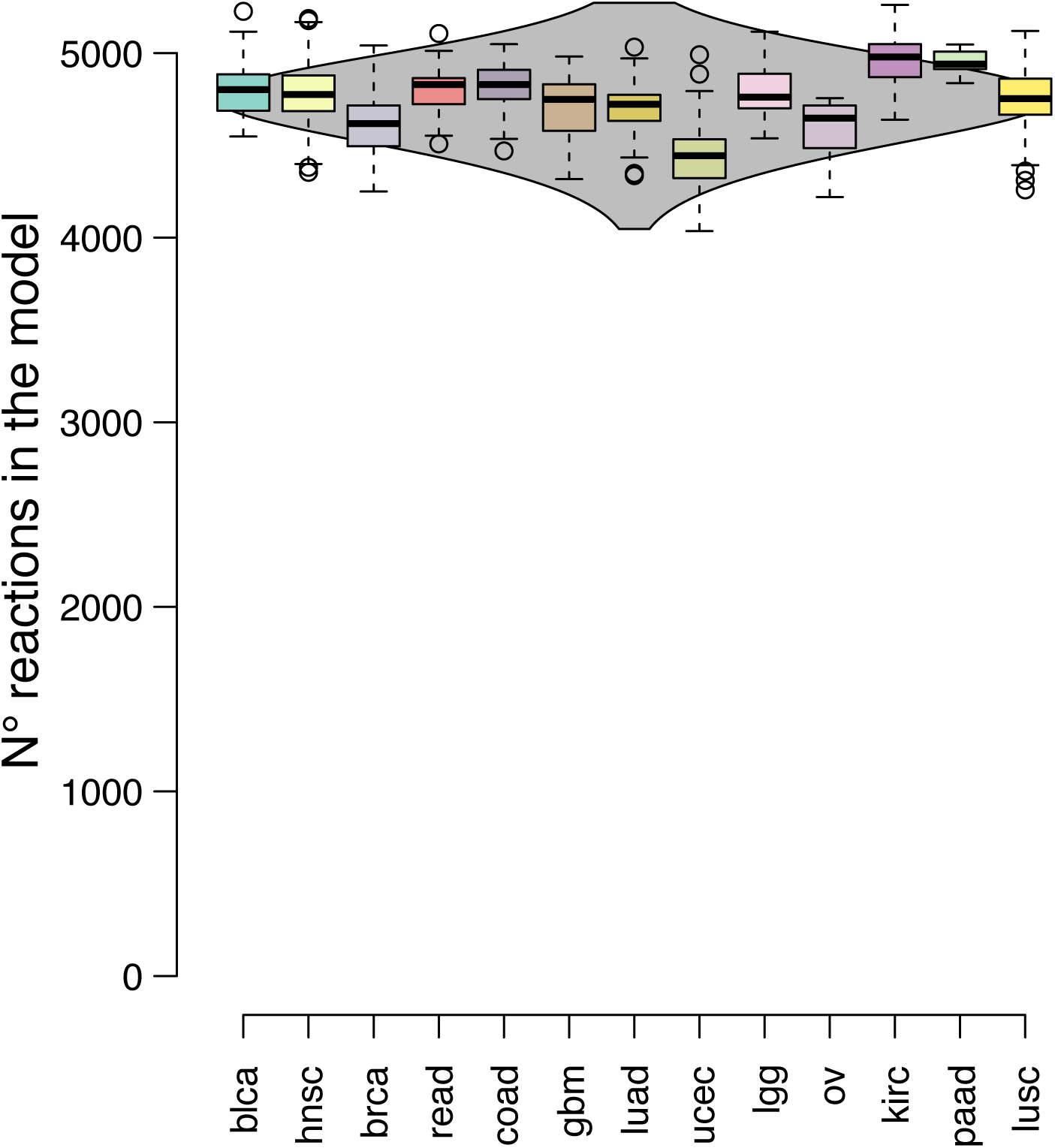
Density boxplot of the number of reactions included in all models (grey), sub-grouped by cancer type. Key as in Fig. S2.

**Figure S5.**
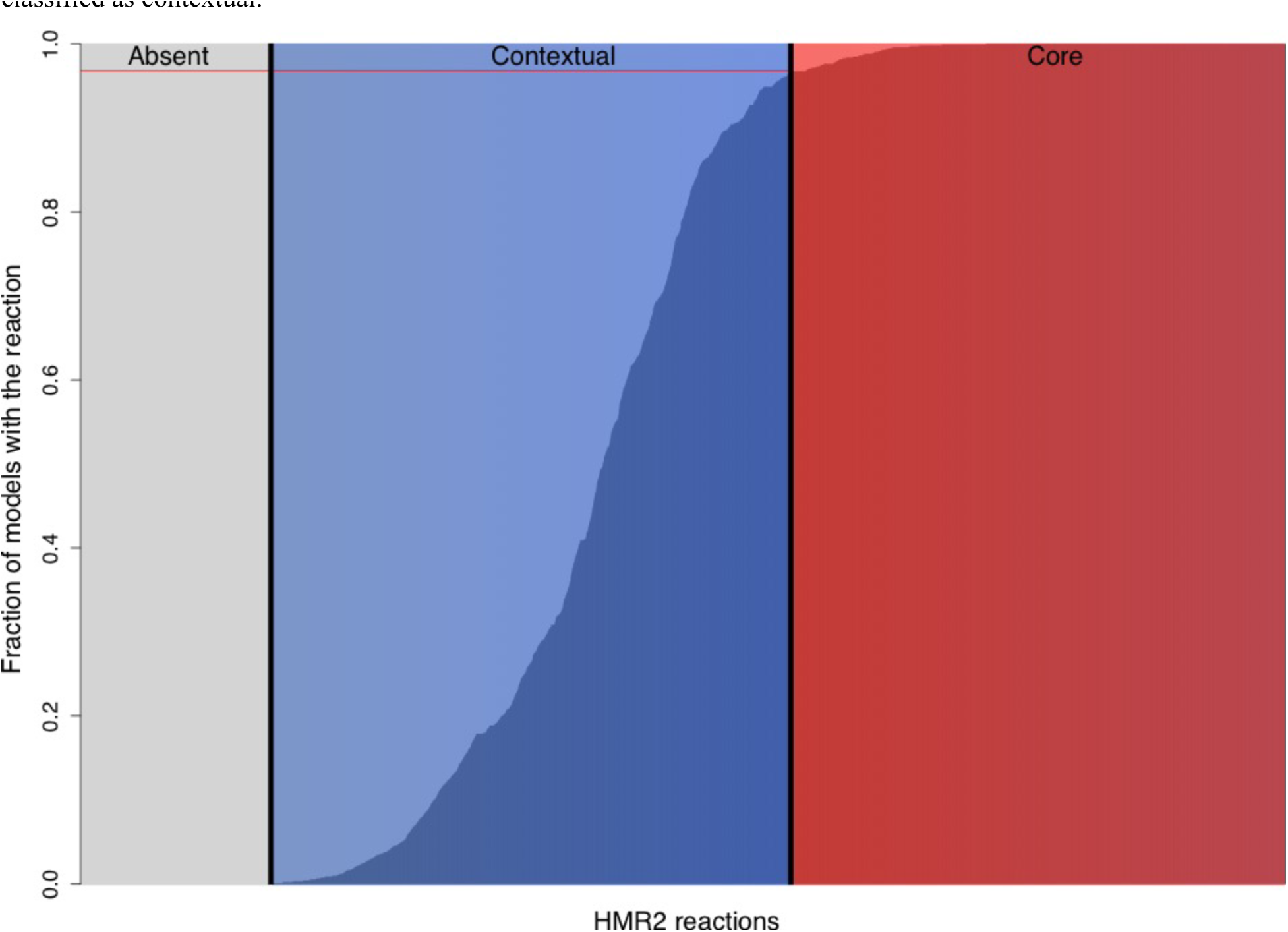
Fraction of models that include a given reaction from the reference model *HMR2* (bars). If a reaction is included in more than 95% of models (872 out of 917 models, red horizontal line), it is classified as a core reaction. If excluded in more than 95% of the model, then it is classified as an absent reaction. Any reaction in between is classified as contextual.

**Figure S6.**
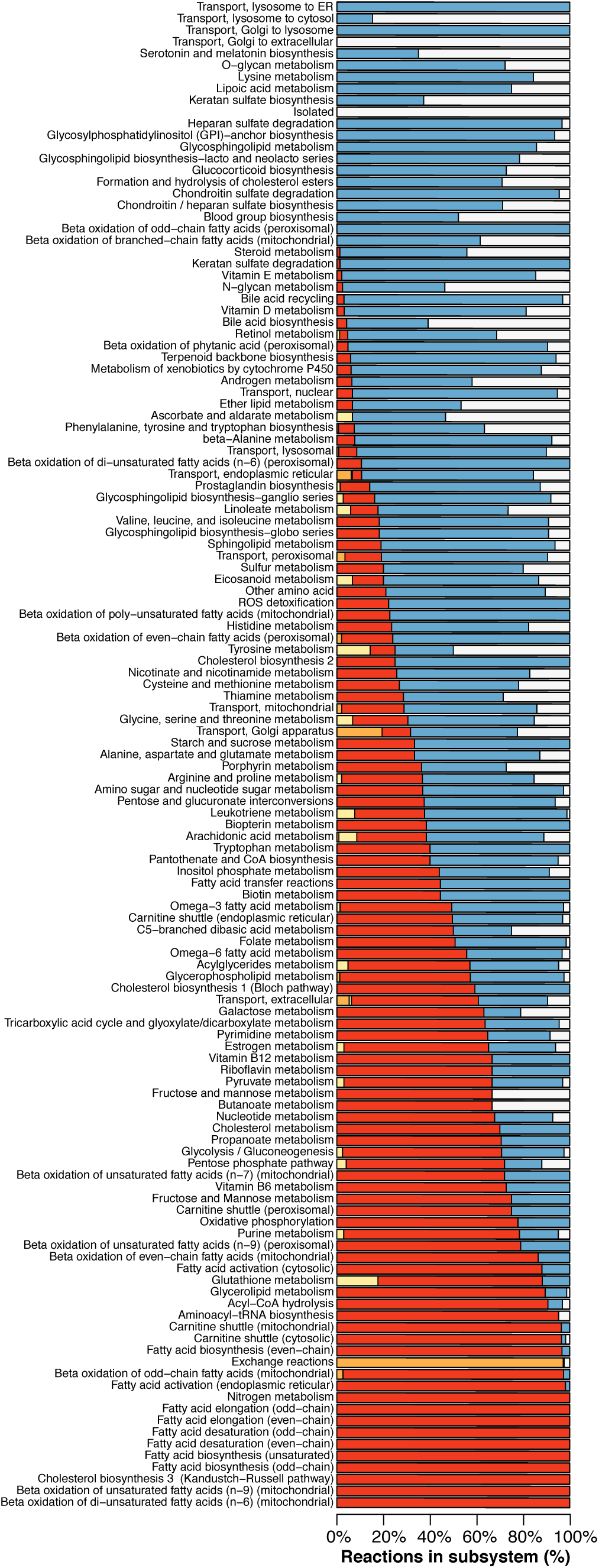
Classification of reactions in the different metabolic sub-systems (same color code as used in Fig. 2B).

**Figure S7.**
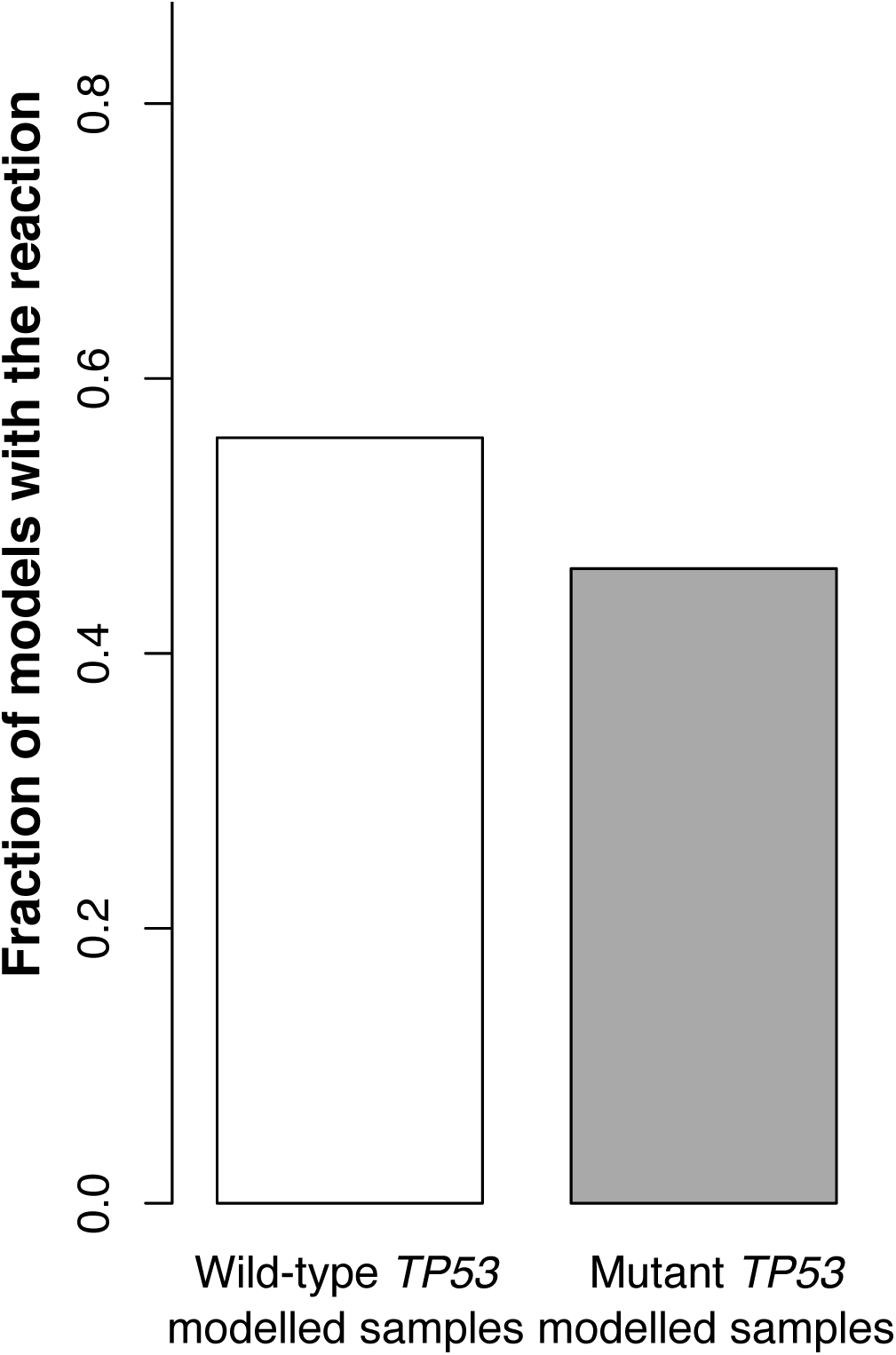
Glutamine conversion to glutamate by glutaminase 2 (GLS2) is the only contextual reaction with a significant association with a mutation, *TP53*. In the barplot, the fraction of models derived from samples with wild-type *TP53* vs. mutated *TP53* that include the *GLS2*-encoded reaction.

**Figure S8.**
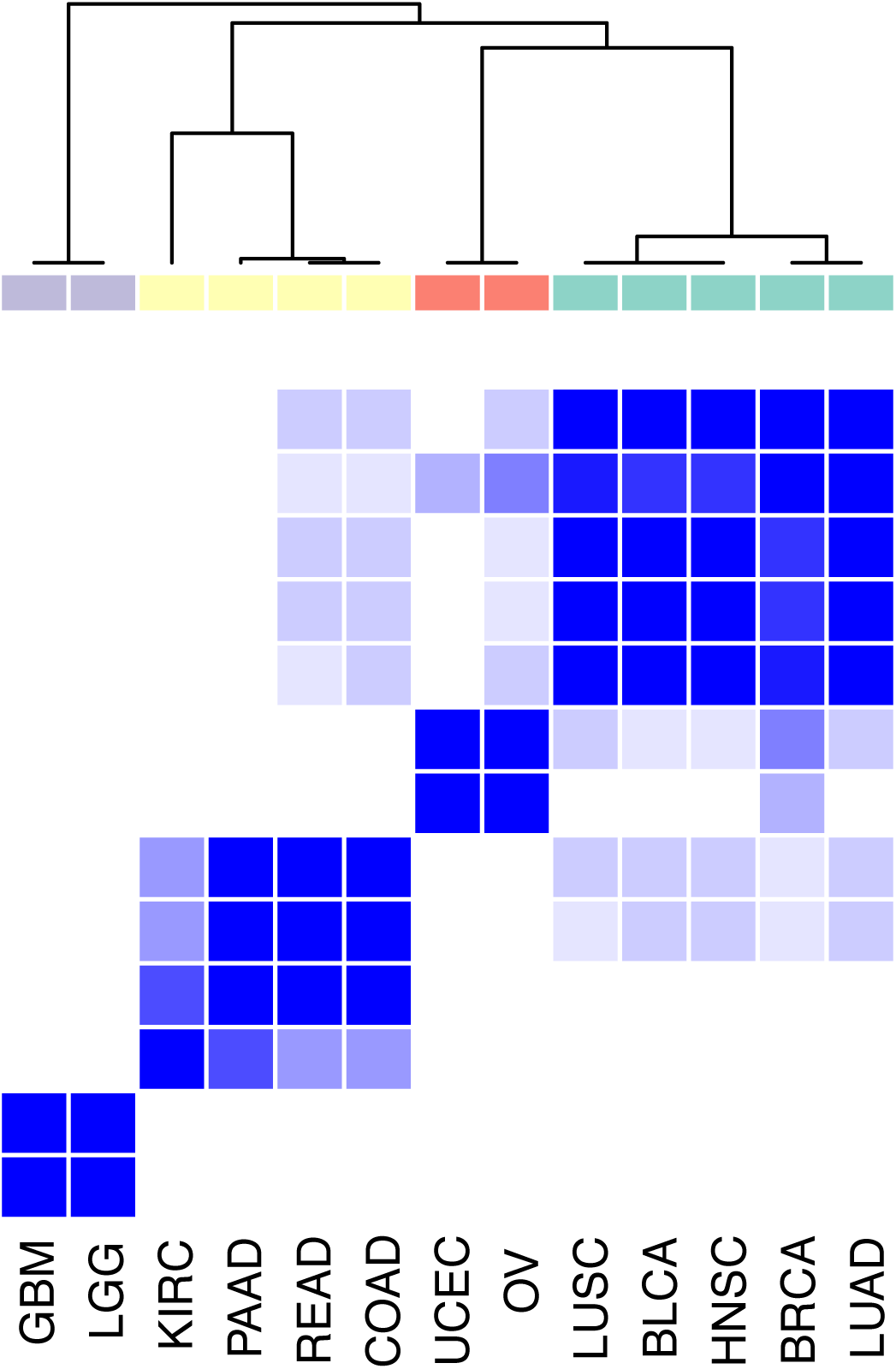
Consensus hierarchical clustering of the 3,269 contextual reactions into cancer types based on the fraction of models representing a cancer type that contain that reaction. Four consensus clusters emerged for cancer type with similar inclusion of contextual reactions.

**Figure S9.**
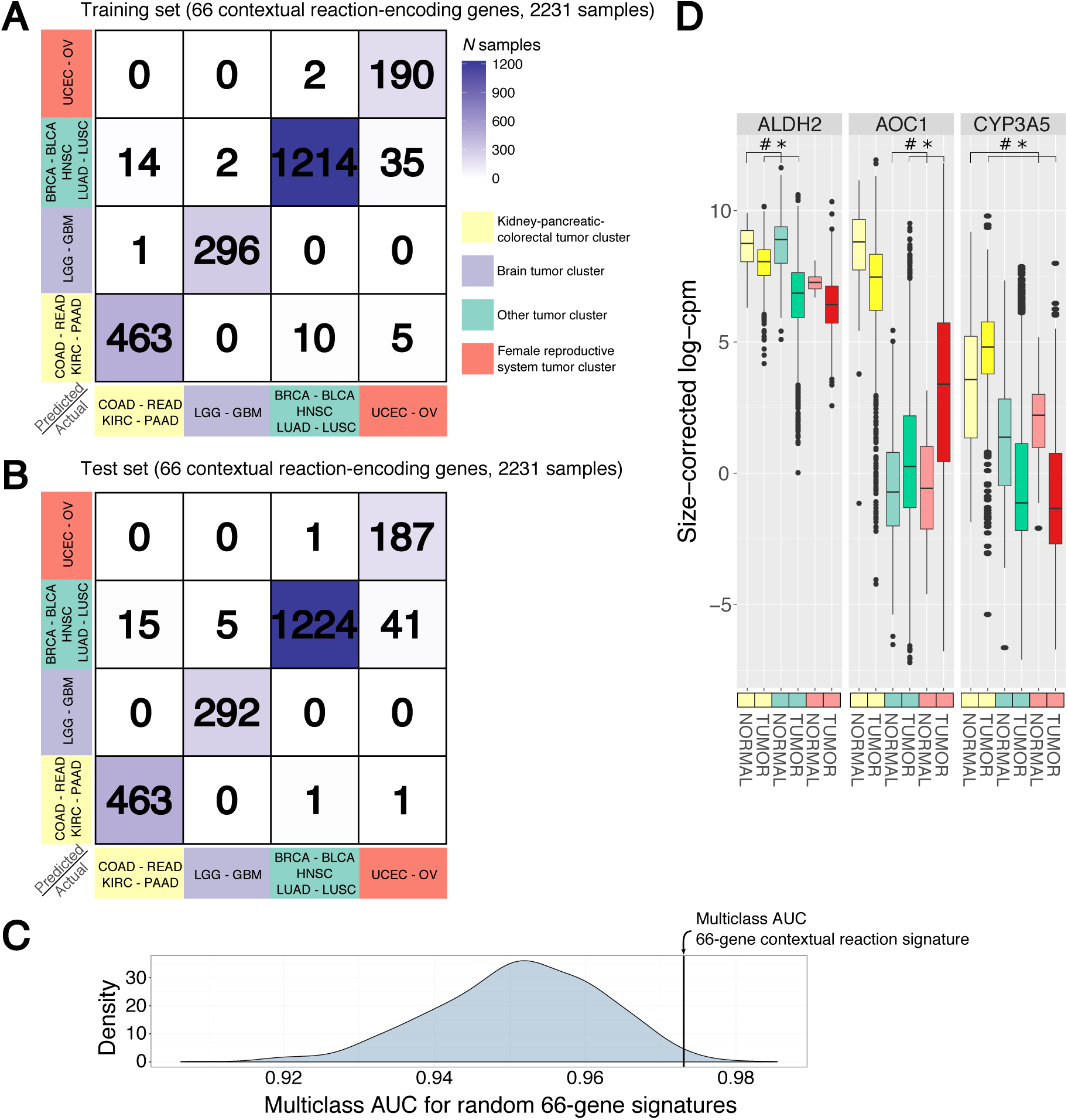
Validation of contextual reaction specificity to the different cancer type clusters-A) Confusion matrix for the training set of a random forest classifier based on the expression level of 66 contextual reaction-encoding genes in an independent group of 2,231 tumor samples. B) Confusion matrix for the test set (2,231 samples) of the classifier trained in A). C) Performance (measured in multiclass AUC) of random forest classifier trained and tested on the same samples but constructed upon 1,000 random signatures of 66 genes. The multiclass AUC for the signature based on the 66 contextual reaction-encoding genes is shown by the vertical line. D) Expression of representative contextual reaction-encoding genes in tumor vs. normal samples grouped by cancer type cluster (3,867 tumor vs. 438 normal samples). Key: *, FDR < 10^−3^; #, FDR > 0.01.

